# Ultra-low frequency neural entrainment to pain

**DOI:** 10.1101/759233

**Authors:** Y Guo, RJ Bufacchi, G Novembre, M Kilintari, M Moayedi, L Hu, GD Iannetti

**Affiliations:** Department of Neuroscience, Physiology and Pharmacology, University College London, London, UK; Neuroscience and Behaviour Laboratory, Istituto Italiano di Tecnologia, Rome, Italy; Faculty of Dentistry, University of Toronto, Toronto, Canada; CAS Key Laboratory of Mental Health, Institute of Psychology, Chinese Academy of Sciences, Beijing, China; Department of Psychology, University of Chinese Academy of Sciences, Beijing, China

**Author notes:** Correspondence to: GD Iannetti, Department of Neuroscience, Physiology and Pharmacology, University College London. Gower Street, London, WC1E 6BT.

## Abstract

Nervous systems exploit regularities in the sensory environment to predict sensory input and adjust behavior, and thereby maximize fitness. Entrainment of neural oscillations allows retaining temporal regularities of sensory information, a prerequisite for prediction. Entrainment has been extensively described at the frequencies of periodic inputs most commonly present in visual and auditory landscapes (e.g. >1 Hz). An open question is whether neural entrainment also occurs for regularities at much longer timescales. Here we exploited the fact that the temporal dynamics of thermal stimuli in natural environment can unfold very slowly. We show that ultra-low frequency neural oscillations preserved a long-lasting trace of sensory information through neural entrainment to periodic thermo-nociceptive input as low as 0.1 Hz. Importantly, revealing the functional significance of this phenomenon, both power and phase of the entrainment predicted individual pain sensitivity. In contrast, periodic auditory input at the same ultra-low frequency did not entrain ultra-low frequency oscillations. These results demonstrate that a functionally-significant neural entrainment can occur at temporal scales far longer than those commonly explored. The non-supramodal nature of our results suggests that ultra-low frequency entrainment might be tuned to the temporal scale of the statistical regularities characteristic of different sensory modalities.

## Introduction

The sensory environment is dynamic in nature, with its temporal structures unfolding across multiple timescales. Time is therefore an indispensable aspect of sensory experiences. The ability of the brain to track and predict the temporal dynamics of sensory inputs allows an organism to take appropriate actions to meet the changing environmental demands. What neural processes may be responsible for these functions, however, is still an open question. Accumulating evidence supports a theory that neural codes of temporal information build on brain oscillations (1–3). Taking situations involving rhythmic sensory inputs as an example, the brain may adapt to the external rhythm through entrainment of ongoing neural oscillations at the corresponding frequency (2, 4–6). Thus, neural entrainment constitutes a flexible mechanism through which the brain adjusts the power or the phase of ongoing oscillations as a function of sensory input, with consequences on brain dynamics that can persist after the sensory input has ceased (7, 8).

Temporal dynamics are ubiquitous in sensory domains, including pain (9–13). Despite this fact, most neuroimaging studies investigating the neural mechanisms of pain were conducted using transient painful stimulation (14–16). This approach poses two main problems. First, the neural responses to transient painful stimulation are dominated by supramodal neural activities - i.e. activities associated with the detection of salient environmental events regardless of their modality (17–19) - which limits the usefulness of this approach in identifying nociceptive-specific brain processing (16). Second, the presentation of a single intense painful stimulus does not capture the rich and often long-lasting dynamics of pain perception, leaving the critical question of how the brain processes dynamic pain information largely unanswered. There is, therefore, a growing consensus in the field that a shift is needed from measuring brain responses elicited by transient painful stimulation to more naturalistic approaches that allow the capture of the temporal dynamics of pain (15, 16).

Some attempts have been made in this new direction (10, 20–25). Tonic and fluctuating nociceptive stimuli delivered at temporal scales of seconds to minutes have been used to better simulate the dynamics of spontaneous pain in clinical conditions (14, 15). A small number of studies tried to relate the temporal profile of brain activity sampled using electroencephalography (EEG) to that of either nociceptive input or reported pain. The main observations were a relationship between the power of alpha (8-12 Hz) and beta (13-30 Hz) oscillations in sensorimotor areas and the fluctuations of nociceptive input (21, 22, 25), as well as between the power of gamma oscillations (>30 Hz) in medial prefrontal cortex and fluctuations of pain intensity (22, 25). In other words, the slow temporal dynamics of both nociceptive input and pain seem to be reflected in neural oscillations at frequencies several orders of magnitude higher than that of ongoing pain.

The above considerations lead to an outstanding question about the role of low-frequency neural oscillations: can the slow dynamics of pain be represented in neural oscillations at the same timescales? This is physiologically plausible, for several reasons. First, a 1:1 phase locking between the rhythm of external sensory inputs and neural oscillations is theoretically possible (26–28), and has been in fact repeatedly observed in auditory and visual domains in the form of neural entrainment (7, 29–33). In these cases, the neural oscillations, and specifically their phase structures, may serve as a substrate of temporal representation and prediction of the incoming input (1, 2). Second, even when the power of higher frequency oscillations is modulated at the lower frequency of the sensory input, oscillations at the same frequency of the input can still be functionally important: they may coordinate rapidly-changing and local neural processing, usually reflected by high-frequency brain activities, with brain activities operating at slower and more behaviorally-relevant timescales (1, 34, 35).

Therefore, in the current study, we investigated whether the brain encodes long-timescale dynamics of painful input through entrainment of ultra-low frequency neural oscillations. Specifically, we investigated (i) whether and how amplitude and phase modulations contribute to such entrainment, (ii) whether this entrainment is supramodal, and (iii) whether the strength of entrainment reflects the variability of perceptual sensitivity across participants.

To address these questions, we recorded high-density EEG (128 electrodes) from participants receiving continuous thermo-nociceptive and continuous loud auditory stimuli oscillating at 0.1 Hz. Painful stimuli were delivered over the hand dorsum using a feedback-controlled laser stimulator with high temporal and temperature precision. Participants were requested to rate the perceived stimulus intensity on a visual analogue scale (VAS). To control for the confounding effect of the rating task, in two additional conditions participants received painful and auditory stimuli but were not required to rate them. Finally, to control for stimulus-intensity effects, we included an additional condition in which painful stimuli of lower intensity were delivered. Thus, there were five conditions in total.

Our results provide clear evidence that the long-timescale dynamics of nociceptive input are encoded by neural oscillations at the same ultra-low frequency of the input. The observations that this encoding was dependent on the phase of ongoing neural oscillations, and particularly that it outlasted the stimulus input, imply a true neural entrainment mechanism. This ultra-low frequency entrainment was not supramodal, as it was robust during nociceptive stimulation but not present during auditory stimulation of similar intensity. Remarkably, the strength of neural entrainment to the nociceptive input was predictive of pain sensitivity across individuals.

## Results

### Stimulus input and perceptual ratings have similar temporal profiles

Participants’ continuous rating of perceived intensity roughly followed the temporal pattern of the rhythmic stimulation. In both pain (**Fig 1A**) and auditory (**Fig 1B**) rating time series, we observed three cycles whose period was similar to that of the stimulus. We formally compared the peak amplitude and latency of ratings across conditions and cycles. The results (**Fig 1C** and **Supplementary Information**) showed that auditory ratings peaked earlier than pain ratings in all three cycles, and that pain ratings were decreased and delayed in the last two cycles compared to the first one (**Fig 1C**).

**Figure 1.**
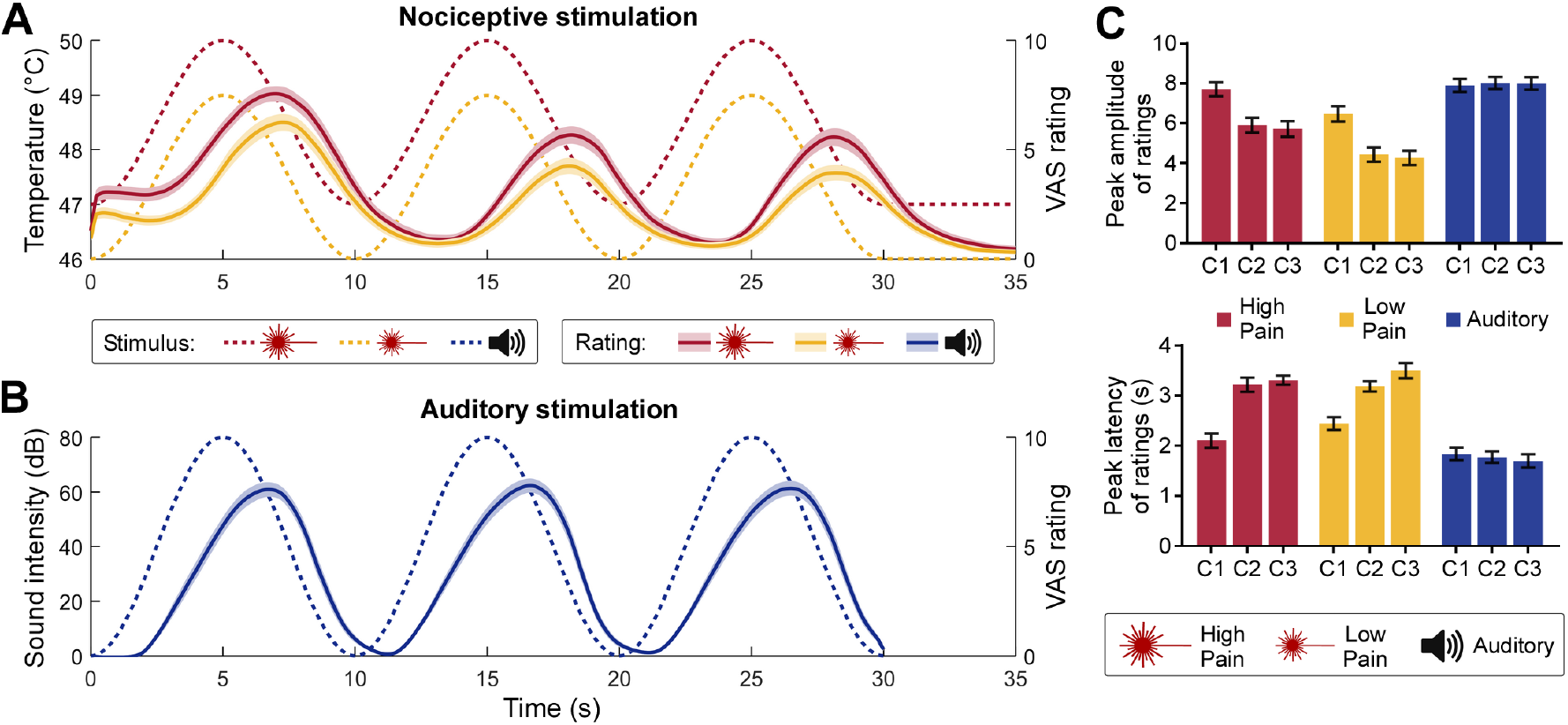
Rhythmic stimulus inputs and perceptual ratings. (**A**) Nociceptive stimuli consisted of a 0.1-Hz sinusoidal modulation of skin temperature with a 3°C difference between peaks and troughs, lasting 30 s. Stimulation temperatures were adjusted for each subject (see Materials and Methods). High-pain stimuli (dotted red line) were set to 1°C above the low-pain stimuli (dotted yellow line). In some conditions, participants (N=30) were required to continuously rate their perceived pain intensity using a visual analogue scale (VAS) ranging from 0 to 10. Both the high-pain (solid red line) and low-pain (solid yellow line) rating time courses followed the nociceptive input (shaded regions indicate the standard error of the mean [SEM] across subjects). (**B**) Auditory stimuli consisted of a 0.1-Hz sinusoidal modulation of the amplitude of a pure tone at 280 Hz (dotted blue line). In some conditions participants were required to continuously rate their perceived sound intensity (solid blue line). (**C**) Peak amplitude (top) and latency (bottom) of the intensity ratings. The peak latency is expressed as difference between a peak in the rating and the corresponding peak in the stimulus. C1-C3: cycle 1 to 3. Error bars indicate the SEM across subjects.

### Low-frequency nociceptive input enhances neural activity at the frequency of the stimulus

Next, we examined whether brain activities encoded the low-frequency rhythmic stimulation. When inspecting the time-domain EEG responses to the nociceptive input, we observed a clear oscillatory pattern reminiscent of that of the stimulus at central electrodes (**Fig 2A**). Importantly, these EEG oscillations were not visible in response to the auditory stimulation. To quantify the frequency contents of these EEG responses, we transformed single-trial EEG signals into the frequency domain and then averaged the resultant power spectra across trials, for each subject and condition (**Fig S1**). Note that we performed the frequency decomposition at trial level rather than on single-subject average waveforms because in the former case it is possible to detect an increase of power regardless of whether neural activity is phase-locked across trials. To reveal the frequency of power increases, we subtracted the average power of neighboring frequencies from each given frequency, yielding background-subtracted power (BSP), as previously recommended (20, 36, 37). We found strong evidence for a power enhancement only at 0.1 Hz for all three conditions entailing nociceptive stimulation (**Fig 2B;** all *t*_29_>7.802, *P*<0.0001, Cohen’s *d*>1.424). This effect was maximal in central scalp regions (**Fig 2B**; also see **Fig S2** for detailed results at other frequencies and scalp positions). This power enhancement could be consistently detected across single trials in the majority of subjects (**Fig 2D**).

**Figure 2.**
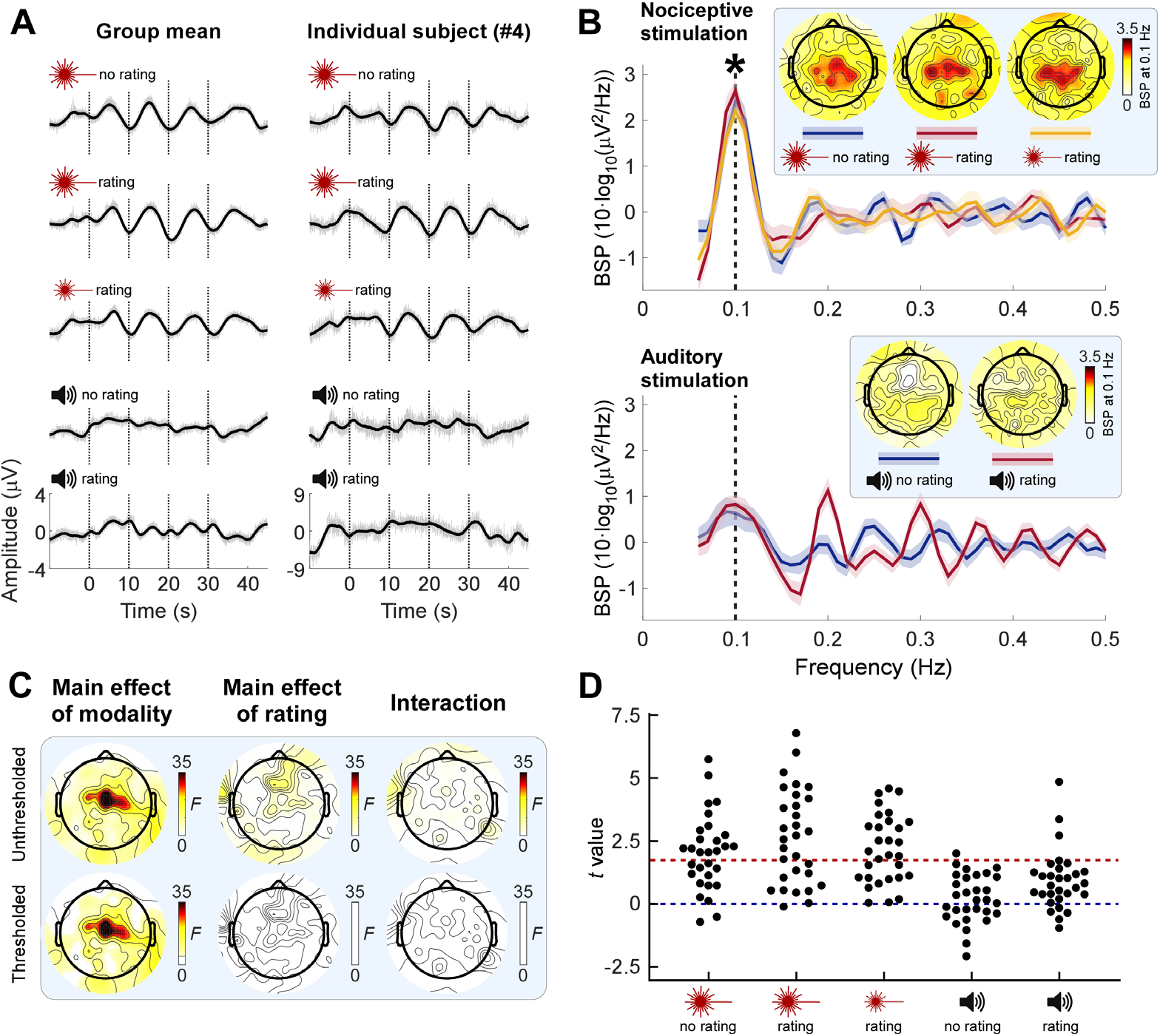
Neural oscillations are enhanced at the frequency of nociceptive input. (**A**) EEG time series averaged across subjects (left column; N=30) and from a representative subject (right column) in a central electrode cluster (Cz and its closest neighbours FCC1h, FCC2h, CCP1h, and CCP2h). Note how the neural activity displayed an oscillatory pattern reminiscent of that of sensory input in pain, but not in auditory conditions. Signals smoothed with a moving mean filter with a 2-s window (black) are superimposed on unsmoothed signals (gray). (**B**) Background-subtracted power (BSP) of EEG in the central electrode cluster during nociceptive (top) and auditory (bottom) stimulation. Shaded regions around the solid lines indicate SEM across subjects. Consistent power enhancement across subjects (marked by asterisk; one-sample *t*-test of BSP against 0, FDR corrected across frequencies) was only observed at the stimulus frequency (0.1 Hz, vertical dashed line) during nociceptive stimulation. Insets show the scalp topographies of the BSP at 0.1 Hz. (**C**) Topographies of *F* values from a two-way repeated-measures ANOVA exploring the effect of factors Modality and Rating on 0.1-Hz BSP. Top row: *F* values; bottom row: thresholded *F* values (*P*<0.05, FDR corrected across electrodes); non-significant regions are masked with white. (**D**) Single-subject *t* values expressing the across-trial consistency of 0.1-Hz power enhancement at central electrodes. Red dashed line: *t* value at *P*=0.05.

We observed strong evidence that the BSP at 0.1 Hz in central scalp regions was greater in pain than in the auditory conditions (main effect of Modality: *F*_1,29_=39.48, *P*<0.0001, partial *η*^2^=0.5765, two-way repeated-measures ANOVA at Cz; see **Fig 2C** for scalp topography of this effect). There was no evidence for a main effect of Rating, nor for a Modality x Rating interaction (**Fig 2C**), indicating that the power enhancement was not dependent on the rating task.

These results showed an enhancement of neural activities specifically at the frequency of rhythmic painful stimulation. Importantly, this effect was neither supramodal (i.e., it was not present in audition), nor dependent on whether the participants were performing a rating task. Also, the observation of an increase of BSP calculated using an extremely narrow frequency window of 0.06 Hz (i.e. from −0.03 Hz to +0.03 Hz with respect to 0.1 Hz) strongly indicates that the observed power increase is not consequent to a general autonomic response, but has instead a neural origin.

### Phase reorganization of neural activity at the frequency of the nociceptive stimulus

The finding that neural activity and stimulus profile have the same peak frequency does not necessarily indicate a stable phase relationship between the two (38). To test for such a phase relationship, we quantified the EEG phase locking across trials using inter-trial phase coherence (ITPC), separately for each subject and condition. Since EEG trials were time aligned to the onset of the rhythmic stimulation, it follows that here ITPC also quantified the consistency of the phase relationship between stimulus and EEG data within each individual. We observed that during painful, but not auditory, stimulation there was a clear peak of ITPC at 0.1 Hz. This was observed in both the mean ITPC across subjects (**Fig 3A**) and the percentage of subjects with significant ITPC (**Fig 3B**; ITPC significance was determined in each subject using the Rayleigh’s test for circular uniformity (39)). This phase-locking effect at the frequency of the nociceptive stimulation was maximal in central scalp regions (**Fig 3A-B**). To test for the group-level consistency of this effect, we compared the group-mean ITPC and the percentage of subjects with significant ITPC to randomized data (see Materials and Methods). In all pain conditions, both ITPC measures at 0.1 Hz in central regions were consistently greater than chance (all *P*<0.001) (**Fig 3A-B**; also see **Fig S3** for detailed test results at other frequencies and positions).

**Figure 3.**
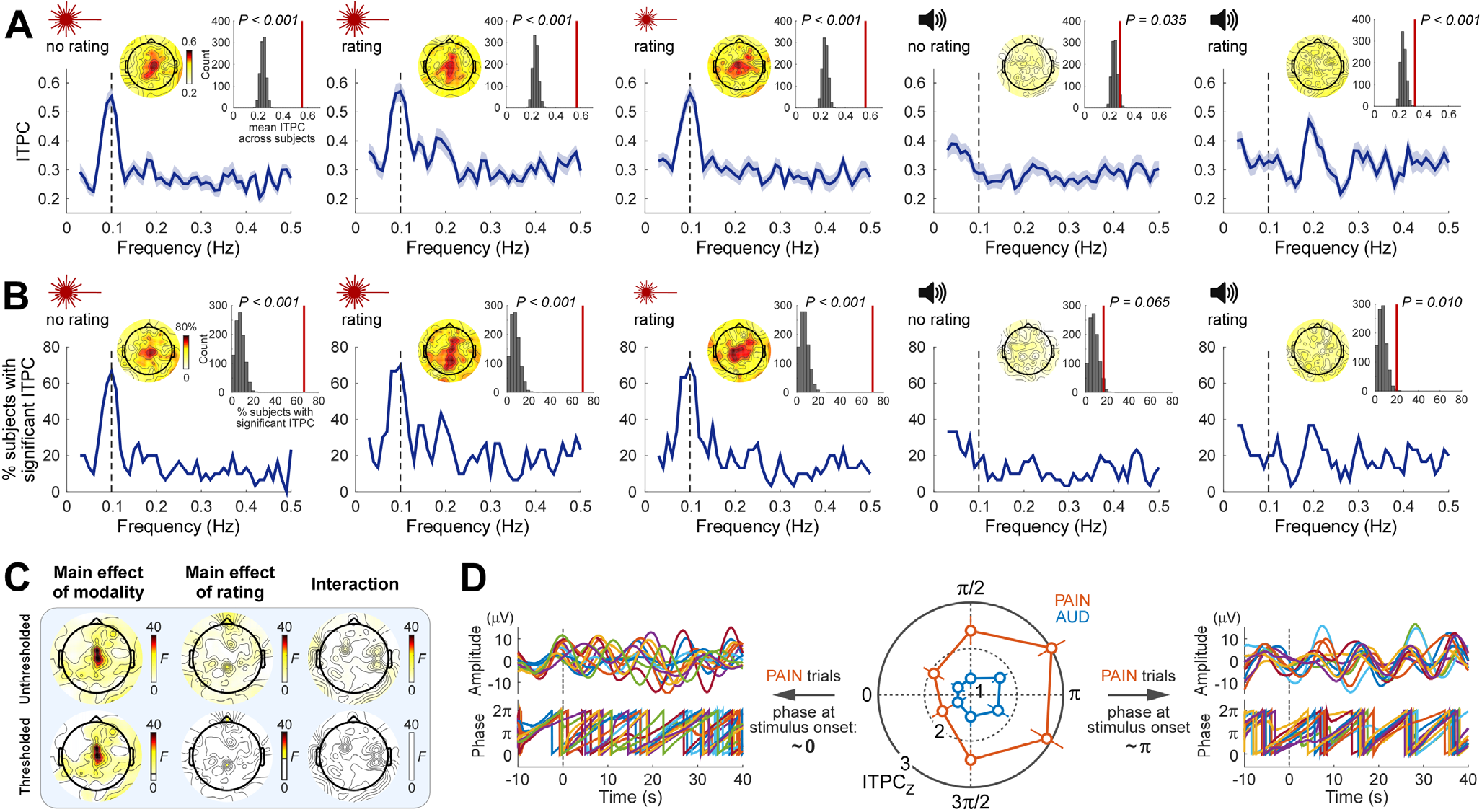
Rhythmic nociceptive input adjusts the phase of neural oscillations at stimulus frequency. (**A**) Inter-trial phase coherence (ITPC) averaged across subjects (N=30) as a function of frequency at central electrodes (blue lines; shaded regions indicate SEM across subjects). Note that pain conditions show an ITPC peak at the stimulus frequency (0.1 Hz, vertical dashed line), whereas the auditory conditions do not. Insets show topographies of the ITPC at 0.1 Hz and the comparison of the 0.1-Hz ITPC from central electrodes (vertical red lines) to ITPC obtained from randomized data. (**B**) Percentage of subjects with significant ITPC (Rayleigh’s test *P*<0.05) as a function of frequency at central electrodes. Note the peak at 0.1 Hz during nociceptive but not auditory stimulation. Insets show topographies of the percentage of subjects with a significant ITPC at 0.1 Hz and the comparison at central electrodes to the same percentage obtained from randomized data. (**C**) Topographies of *F* values from a two-way repeated-measures ANOVA exploring the effect of factors Modality and Rating on 0.1-Hz ITPC. Top row: *F* values; bottom row: thresholded *F* values (*P*<0.05, FDR corrected across electrodes); non-significant regions are masked with white. (**D**) The degree of phase locking during nociceptive stimulation was dependent on the phase of prestimulus oscillations. The strongest phase locking occurred in trials in which the stimulus started around the trough (π rad) of ongoing oscillations (right: pain trials in the two bins around π from a representative subject), whereas the weakest phase locking occurred when the stimulus started around the peak (0 rad) of ongoing oscillations (left: pain trials in the two bins around 0 from the same subject). Error bars indicate SEM across subjects.

We observed strong evidence that the 0.1-Hz ITPC in central scalp regions was higher in pain than in audition (main effect of Modality: *F*_1,29_=40.15, *P*<0.0001, partial *η*^2^=0.5806, two-way repeated-measures ANOVA at Cz; see **Fig 3C** for scalp topography of this effect). There was no main effect of Rating on 0.1-Hz ITPC, except in two electrodes distant from each other (Fpz, CPz), in which the evidence for this main effect was, however, weak (**Fig 3C** for scalp topography of this effect). There was no Modality x Rating interaction (**Fig 3C**).

Given that phase locking across trials (i.e., a relatively stable phase relationship between the neural responses and the stimulus profile) could result from a reorganization of the phase of ongoing neural oscillations, we tested whether the degree of phase locking was related to the phase of ongoing EEG. We sorted single trials into six bins according to the prestimulus phase at central scalp regions and calculated 0.1-Hz phase locking (ITPC_Z_; Rayleigh’s Z, see Materials and Methods) during stimulation for each bin, modality, and subject. As shown in **Fig 3D**, we observed clear evidence that the phase of prestimulus oscillations influenced the degree of phase locking during stimulation. Phase locking in the pain trials was maximal when the onset of the rhythmic nociceptive stimulation coincided with the trough of ongoing oscillations (i.e., around π) and minimal when the stimulus onset coincided with the peak (i.e., around 0 or 2π) (*F*_5,145_=4.433, *P*=0.0009, partial *η*^2^=0.1326, two-way repeated-measures ANOVA, main effect of Bin; post hoc tests showed significant differences between either of the two bins around π and either of the two bins around 0, all *P*<0.0054). Although there was strong evidence that the phase locking in auditory trials was lower than that in pain trials (*F*_1,29_=18.31, *P*=0.0002, partial *η*^2^=0.3870, main effect of Modality), the lack of a Bin x Modality interaction (*F*_5,145_=1.018, *P*=0.4094, partial *η*^2^=0.0339) indicated that also in the auditory trials phase locking was influenced by the phase of prestimulus oscillations, although to a lesser extent (there were no differences in phase locking between bins in auditory trials; *P*>0.1440 in all bin comparisons). This dependence of phase locking on the phase of prestimulus activities indicates an entrainment of intrinsic neural oscillations to the rhythm of nociceptive stimulation rather than just the occurrence of stimulus-evoked responses (for detailed discussions see, e.g., (40, 41)).

Altogether, these results show phase locking of neural oscillation at the frequency of the rhythmic nociceptive stimulation. The degree of such phase locking clearly depended on the phase of ongoing neural oscillations. Similar to the power increase, this phase-locking effect was also not supramodal, and not related to whether the participants were performing a rating task.

### Pain sensitivity across individuals is reflected in the strength of neural entrainment

Perceived pain was stronger in participants with a stronger neural entrainment. Specifically, we correlated the pain rating time series to the indices of power enhancement (BSP) and phase locking (ITPC) at 0.1 Hz in central scalp regions. We observed clear positive correlations in the time interval around each rating peak (**Fig 4**). Remarkably, the across-subject variability in perceived pain intensity was also reflected in the phase relationship between the entrained oscillations and the stimulus (**Fig 5**). To evaluate this relationship, we fitted a single-cycle cosine to the subject-mean peak rating as a function of the phase of 0.1-Hz oscillation in central scalp regions. In subjects who rated the nociceptive stimulus as more painful the phase of neural activity at 0.1 Hz was closer to that of the stimulus input. This is an important finding given that almost all nociceptive-evoked neural responses fail to track pain sensitivity across subjects, in both human and animal studies (42).

**Figure 4.**
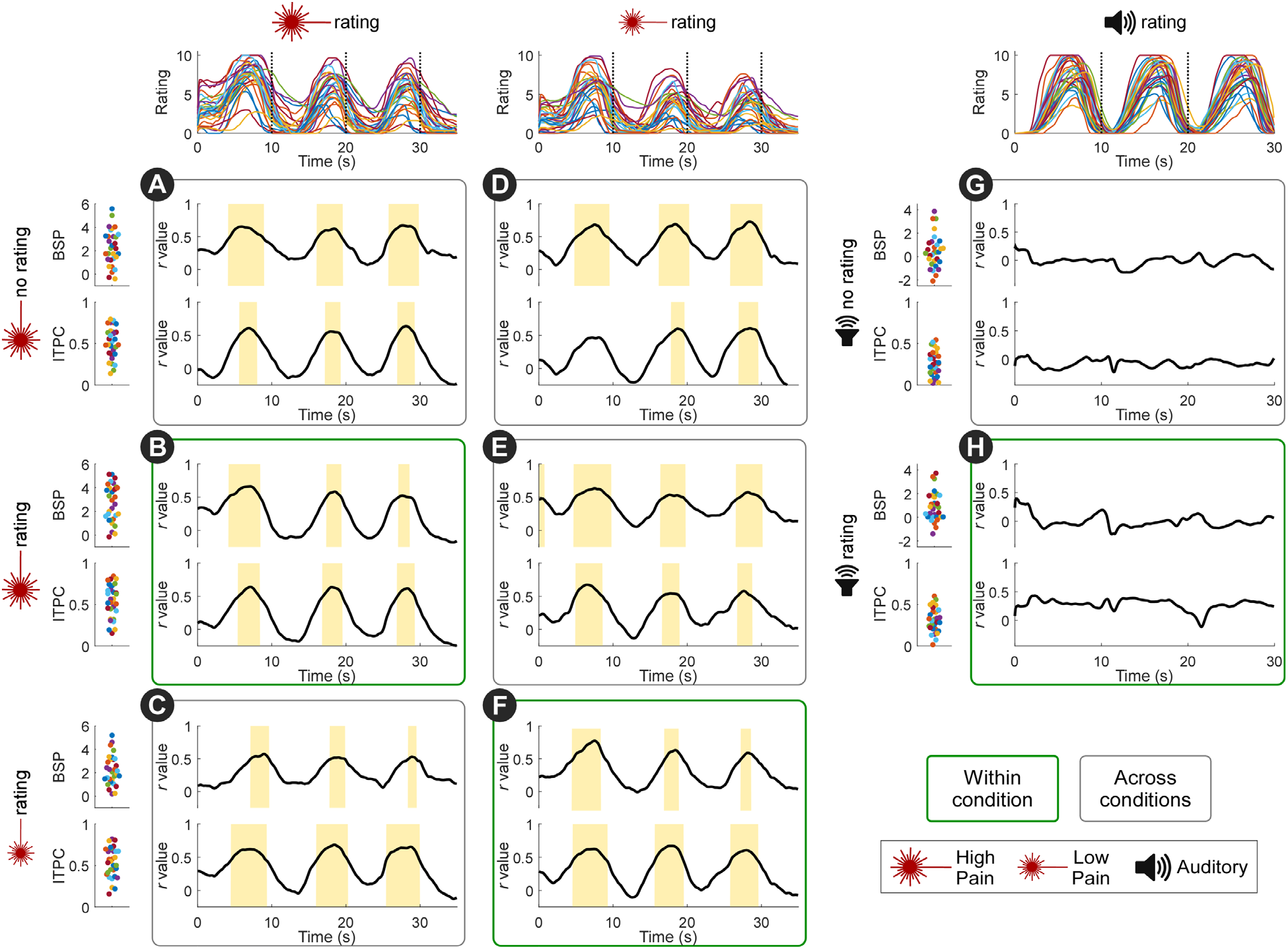
Pain sensitivity across individuals is reflected in the strength of neural entrainment: power and phase locking. Traces on the top show individual rating time series. Scatter plots on the left show individual background-subtracted power (BSP, top scatterplots of each condition, expressed as 10·log_10_(μV^2^/Hz)) and inter-trial phase coherence (ITPC, bottom scatterplots of each condition) at 0.1 Hz at central electrodes. Black lines show the correlation coefficients between BSP/ITPC and ratings across time. Both BSP and ITPC of each of the three pain conditions were correlated to the high pain (**A-C**) and low pain (**D-F**) ratings, in the time intervals around the rating peaks (marked by yellow bars; *P*<0.05, FDR corrected across time points). Note that the positive across-subject correlation was present not only when correlating BSP and ITPC with ratings within condition (**B,F**), but also when correlating BSP and ITPC with ratings across conditions (e.g. when correlating BSP and ITPC from the pain no-rating condition with high-pain and low-pain ratings; **A,D**). Importantly, BSP and ITPC in the auditory conditions were not correlated to the auditory intensity ratings (**G-H**). N=30 subjects.

**Figure 5.**
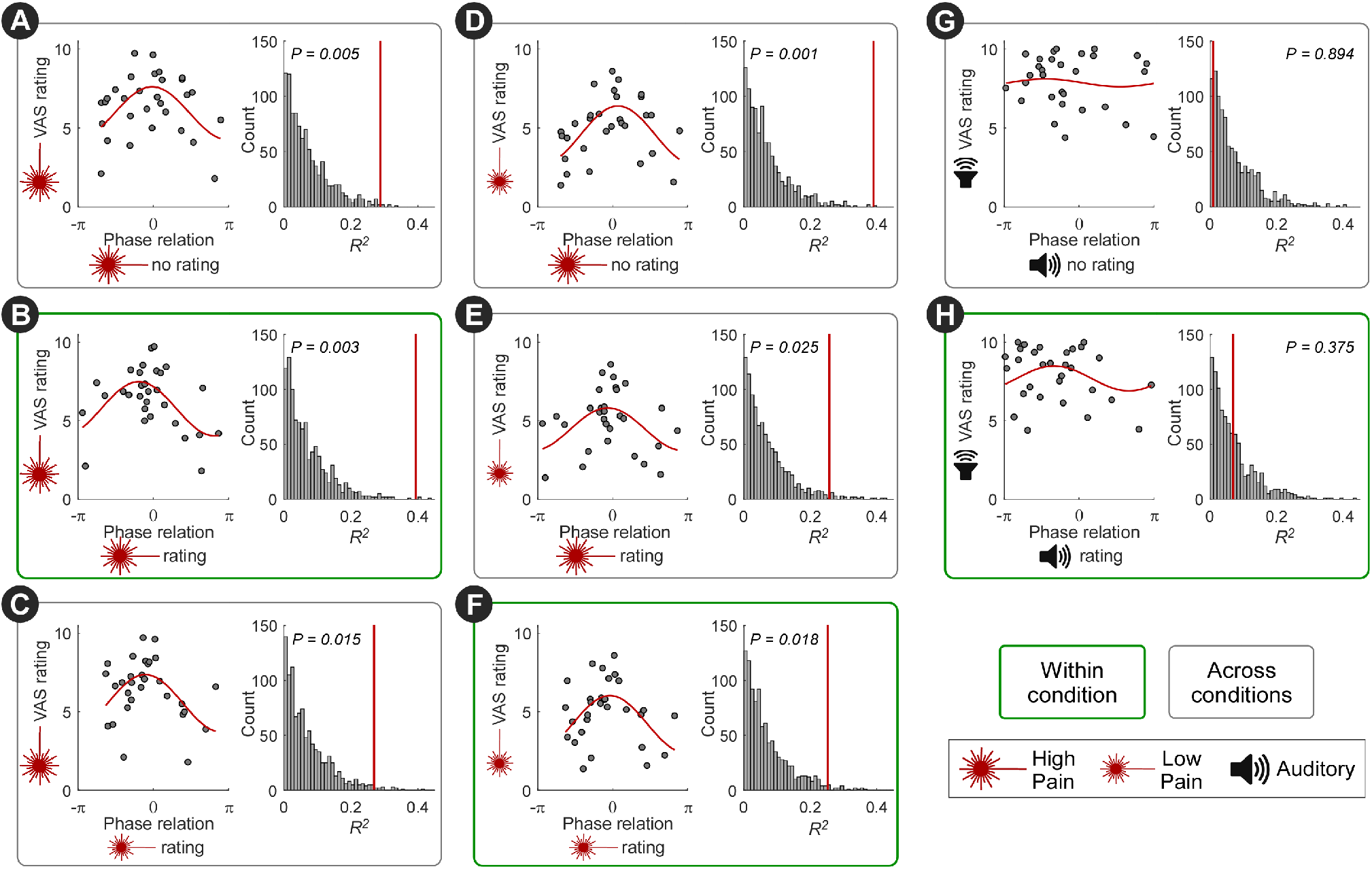
Pain sensitivity across individuals is reflected in the strength of neural entrainment: phase. The across-subjects variability in perceived pain intensity was also reflected in the phase relationship between the entrained oscillations and the stimulus profile. For each subject, the mean of high pain (**A-C**), low pain (**D-F**), or auditory (**G-H**) peak ratings was plotted against the phase difference between the 0.1-Hz oscillation at central electrodes and the stimulus (“phase relation”). To evaluate this relationship, we fitted a single-cycle cosine function (red lines). Coefficient of determination (*R*^2^) of the cosine fit was tested by random permutation of the phase across subjects. Subjects who rated the nociceptive stimulus as more painful entrained more closely to the phase of the nociceptive input (i.e., with a phase relation around 0). This relationship was preserved not only within condition (**B,F**) but also across conditions (**A,C-E**). Importantly, such relationship was not present in the auditory conditions (**G-H**). N=30 subjects.

Importantly, all these relationships (i.e. the correlation between BSP/ITPC and pain ratings, and the relationship between phase and pain ratings) existed not only (i) within the conditions entailing nociceptive stimulation with ratings (**Fig 4B, F**; **Fig 5B, F**), but also (ii) between high pain and low pain conditions (**Fig 4C-E**; **Fig 5C-E**), and even (iii) between pain ratings and the neural entrainment in the pain condition without ratings (**Fig 4A, D**; **Fig 5A, D**). The relationships in (ii) suggested that the strength of neural entrainment reflected individual pain sensitivity. The relationships in (iii) further demonstrated that the link between pain sensitivity and the strength of neural entrainment was not driven by the rating task. No such relationships were observed in the conditions entailing auditory stimulation (**Fig 4G-H**; **Fig 5G-H**).

To test whether the variability in stimulus temperature contributed to the above results, we also tested for an across-subjects relationship between stimulus temperature and pain ratings, as well as between stimulus temperature and neural entrainment. We did not observe such relationships (see **Table S1** for detailed statistical results).

Taken together, these results show that the strength of entrainment could predict how sensitive each participant is to painful stimulations.

### Stimulus-induced neural oscillations outlast nociceptive input

A remarkable observation was that neural oscillations around 0.1 Hz continued for at least 10 seconds after the end of the rhythmic nociceptive input (**Figs 2A** **and** **6**). The observation that the scalp topographies at the peak of the additional cycle resembled those of the previous cycles (**Fig 6**) further indicates self-sustained activity of the same underlying neural process. These observations provide additional evidence for an actual neural entrainment of ongoing EEG activities to the nociceptive input. They also provide further evidence that the observed ultra-low frequency oscillations were not consequent to autonomic responses. Finally, the amplitude of this additional oscillation after the end of rhythmic nociceptive stimulation was correlated with pain ratings across subjects (**Fig 6**).

**Figure 6.**
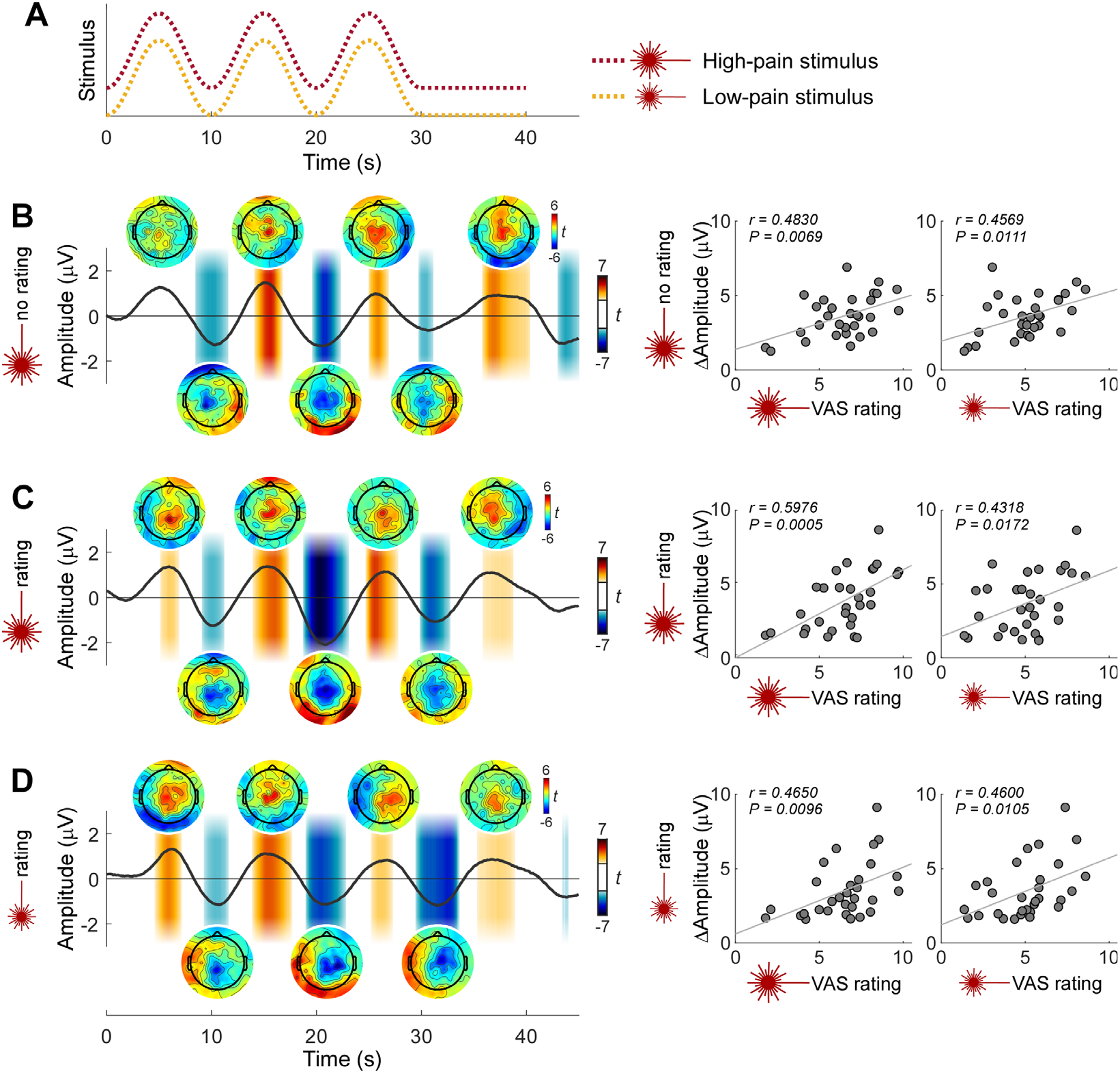
Stimulus-induced neural oscillations outlast the rhythmic nociceptive input. The observation that neural oscillations around 0.1 Hz continued after the end of the rhythmic nociceptive input provides additional evidence for a real neural entrainment of ongoing EEG activity. (**A**) Temporal profile of the high-pain (red dotted line) and low-pain (yellow dotted line) stimulation. (**B-D**) EEG signal at central electrodes was smoothed with a moving mean filter with a 2-s window, linearly detrended, and finally averaged across subjects (N=30) in the conditions entailing high-pain stimulation without rating task (**B**), high-pain stimulation with rating task (**C**), and low-pain stimulation with rating task (**D**). Shaded regions indicate the time windows in which the EEG amplitude is significantly different from 0 (*P*<0.05, point-by-point one-sample *t*-test against 0, FDR corrected across time points). Scalp maps show the *t* value topographies within 1-s window around the peak and trough of each cycle. The similarity of scalp topographies at the peak of the cycle after the end of rhythmic stimulation with those of the previous cycles further indicates self-sustained activity of the same underlying neural process. Plots on the right show that the amplitude difference between the peak and trough of the cycle after the end of rhythmic stimulation was correlated to the mean of the peak pain ratings across subjects.

## Discussion

Here we aimed to identify neural activity present during tonic sensory stimuli that produce slowly fluctuating sensations. We delivered intensity-matched auditory and painful stimuli fluctuating at 0.1 Hz and observed that only painful stimuli resulted in both a power enhancement and a phase locking of brain activity at the same frequency of the stimulus (**Figs 2** **and** **3**). Thus, this brain response was not supramodal, and possibly selective for the somatosensory system. This response likely reflected a true neural entrainment, since (i) the degree of phase locking depended on the phase of ongoing brain oscillations occurring before the onset of the rhythmic input (**Fig 3D**), and (ii) the stimulus-induced brain oscillations outlasted the rhythmic input (**Fig 6**). Importantly, this neural entrainment to the rhythmic painful input was not due to the rating task, as it was also present when participants did not have to rate the painfulness of the stimuli (**Figs 2** **and** **3**). Finally, the strength of the neural entrainment was correlated with pain sensitivity across individuals (**Figs 4** **and** **5**), a relationship that persisted even in the neural activity outlasting the rhythmic stimulus (**Fig 6**).

These findings show that the brain encodes long-timescale dynamics of nociceptive input through entrainment of ultra-low frequency neural oscillations. This work not only represents a step towards analyzing brain processes in more clinically-relevant models of long-lasting and dynamic pain (15, 16), but also sheds new light on the functional significance of neural oscillations at frequencies well below the traditional boundaries of human EEG rhythms.

### Neural entrainment or evoked responses?

The power enhancement (**Fig 2**) and phase-locking (**Fig 3**) of neural activities observed at the frequency of nociceptive input could be explained by either repeated evoked responses or an entrainment of ongoing neural oscillations to the rhythmic stimulus (6, 7, 40, 41, 43–47). Two lines of evidence in our data support an entrainment mechanism.

First, we found clear evidence that the degree of phase locking depended on the phase of ongoing neural oscillations occurring before stimulus onset (**Fig 3D**). Indeed, such phase interactions are predicted by theoretical models and experimental data of entrainment of neural oscillators: although rhythmic stimuli drive brain activities, the effectiveness of this process, reflected in the phase reorganization, also depends on the state of the neural oscillators (33, 38, 48, 49).

Second, as depicted in **Fig 6**, the neural oscillations outlasted the rhythmic painful input. Furthermore, the similarity of scalp topographies of the oscillations during and after rhythmic stimulation suggests common neural generators. This interpretation is corroborated by both theoretical and experimental studies (7, 8, 27, 32, 33, 50–52) demonstrating that entrained neural oscillations can be self-sustaining for a certain amount of time after the end of external rhythm. Future experiments collecting post-stimulus data for longer time intervals will allow a more accurate characterization of the self-sustained activity. Such experiments would also illustrate the recovery process of the oscillatory system from the entrained state.

This neural entrainment reflects processing of nociceptive information that is not captured by more widely-used paradigms employing transient stimuli. Indeed, unlike transient responses that are discrete in time, the entrained oscillations are continuous and might reflect an adaptive internal model of the input temporal regularities, which could facilitate sensory processing of incoming events (1, 2).

### Neural entrainment to rhythmic painful input is independent of rating task

We observed neural entrainment to the rhythmic painful input regardless of whether the participants were required to continuously rate pain intensity (**Figs 2**, **3**, **6**, **S2**, **and** **S3**).

Continuous perceptual ratings are commonly used to investigate the neural correlates of percepts occurring at long timescales (e.g., tonic experimental pain in healthy volunteers (20, 22, 25) or spontaneous fluctuations of pain in chronic pain patients (12, 13)). Such continuous ratings are typically obtained with a finger-span device (12, 13, 22, 25) or a slider (20). While such ratings provide valuable information on the dynamic fluctuations of perception, they heavily confound the analysis of neural data due to the superposition of brain activities related to the motor and cognitive activity related to the rating task (53). A strategy to control for this confound is to ask participants to continuously rate the perceived intensity of stimuli belonging to another sensory modality (e.g., vision) (12, 13, 22, 25, 54): the brain activity measured during painful but not visual stimulation should then reflect pain-selective activity. This paradigm assesses whether the rating task is *sufficient* to explain the brain response sampled during painful stimulation. However, it cannot resolve whether the same brain response remains when no rating task is performed – that is, whether the rating task is *necessary* for observing the brain response. To effectively address this issue, we included in our experiments additional control conditions in which participants had to rate the intensity of neither the auditory nor the painful stimuli.

Thus, our results demonstrate that rating-related brain activities were not *necessary* for the observed entrainment of brain oscillations to the rhythmic painful input. Indeed, the power enhancement, the phase locking, as well as the continuation of the entrained neural oscillations after stimulus offset were also present in the pain condition not involving the rating task (**Figs 2**, **3**, **6**, **S2**, **and** **S3**).

While we did not observe strong evidence for an effect of rating task at the stimulation frequency of 0.1 Hz for either power or phase, the task of rating auditory stimuli enhanced the power and the phase locking at 0.2 and 0.3 Hz (**Figs 2B**, **3A-B** **and** **S3**). In contrast, rating of painful stimuli only slightly enhanced the phase locking at 0.2 and 0.3 Hz (**Fig S3**). This frequency and modality pattern suggests that the rating effect reflects a different mechanism than the neural entrainment to the rhythmic input (44). The fact that the increases and decreases were more regular in auditory than in pain ratings (**Fig 1**) might explain why the effect of rating auditory stimuli was more evident.

### Is the observed neural entrainment modality-specific?

As we discussed above, the entrainment of ultra-low frequency brain oscillations to stimulus input was clearly not supramodal, given that (i) the power increase of EEG signal at stimulus frequency was only consistently observed during nociceptive stimulation but not during loud auditory stimulation (**Fig 2**), and (ii) only the EEG signals during nociceptive stimulation showed a predominant peak of phase locking at stimulus frequency (**Fig 3**). Importantly, these findings by no means imply that the entrainment we observed during nociceptive stimulation was modality-specific. Indeed, strictly speaking, demonstrating the pain specificity of a neural response is virtually impossible, as it would require testing all stimuli that could in principle elicit that response, and show that that response only occurs when pain is experienced (16). Instead, the current findings provide evidence that entrainment at the ultra-low frequency used in this study occurs *preferentially* in response to somatosensory input. That the observed entrainment is preferential to pain would require testing whether it occurs less strongly during non-nociceptive somatosensory stimulation.

The lack of entrainment to the auditory stimuli is not trivial, since there is a considerable amount of evidence for entrainment of neural oscillations to rhythmic auditory stimulation (4, 7, 30, 31, 33, 46). A possible explanation is that the frequency of the delivered auditory stimulation is substantially lower than the timescale of dynamics for which the auditory neural circuits are optimized. Indeed, the temporal structures of speech and music largely occur in the subsecond range (3, 55). Accordingly, previous evidence for auditory entrainment is primarily observed during stimulation at delta (1-4 Hz) and theta (4-8 Hz) frequencies (7, 8, 30, 33, 56, 57).

These previous observations, together with the current results, suggest a different frequency preference for oscillatory entrainment across sensory systems (58). Why would the nociceptive system respond to lower ranges of stimulus frequencies than the auditory system? Given that the temporal dynamics of fluctuations of spontaneous pain and somatosensory detection occur at very low frequencies (11, 59), the brain representations of long-timescale variability might be more essential and relevant to the processing of pain information. These considerations suggest that neural entrainment is tuned to the temporal scale of the statistical regularities characteristic of different sensory modalities, a hypothesis that would require further testing.

### The strength of neural entrainment reflects pain sensitivity across individuals

We observed a clear relationship between perceived pain intensity and the strength of neural entrainment to the nociceptive input. Individuals with higher pain sensitivity had greater power enhancement and phase locking at the frequency of the nociceptive input (**Fig 4**), as well as a phase of the entrained oscillations closer to the phase of the nociceptive input (**Fig 5**). This is a remarkable result, for two reasons. First, even the commonly-observed *within-subject* correlation between nociceptive-evoked responses and subjective pain ratings have been shown to be not obligatory, i.e. they can be easily disrupted by a large number of experimental manipulations (e.g. expectation, stimulus repetition, presentation of non-painful but iso-salient stimuli, congenital insensitivity to pain; (60–62)). Second, almost all nociceptive-evoked neural responses fail to track pain sensitivity *across subjects*, in both human and animal studies (42).

It is important to note that the relationship between entrainment and pain sensitivity across subjects were not driven by the rating task, and were also present when examining conditions entailing different intensities of nociceptive stimulation. Most notably, the relationship between oscillatory amplitude and ratings persisted following the end of rhythmic stimulation (**Fig 6**). Thus, the strength of neural entrainment to rhythmic nociceptive input is, in these experimental conditions, a neural marker of pain sensitivity across individuals.

For clinical purposes, it is important to predict pain across different individuals (63–66). For example, pain may need to be inferred for new patients when verbal reports are unavailable or unreliable, to optimize analgesic treatment and general care (65). Including ultra-low frequency neural entrainment in future pain prediction models may allow for better pain prediction between individuals, especially for the more realistic situations where pain fluctuates as an ongoing percept.

Relevant to this discourse, a recent study revealed that brain oscillations in the gamma band index the variability of pain sensitivity across individuals, in both humans and rodents (42). However, the clinical relevance of this observation is limited because (i) gamma oscillations were elicited by transient nociceptive stimuli causing painful sensations barely reflecting clinical pain, and (ii) due to low signal-to-noise ratio, gamma oscillations are hardly detected in EEG recordings from single individuals. The ultra-low frequency oscillations we described here, in contrast, are (i) elicited by continuous and fluctuating nociceptive input causing ongoing pain that better mimics spontaneous pain in patients, and (ii) can be detected in most single subjects (**Figs 2D** **and** **3B**). Despite these distinctions, these high- and low-frequency indicators of pain sensitivity across individuals could be related. Therefore, an interesting direction for future research is to investigate whether the phase/frequency modulation of slow oscillations and the power modulation of gamma (or other fast) oscillations work synergistically, e.g., to form a hierarchical structure supporting different spatial/temporal scales of brain operation. This could be relevant to tracking and predicting the dynamics of pain, especially in clinical situations (11–13).

## Materials and Methods

### Participants

Thirty healthy human volunteers (18 women; mean age ± SD, 22.8 ± 2.9 years, age range 19–30 years; all right-handed) participated in the study. All participants gave written informed consent and received monetary compensation for their participation. The study was approved by the ethics committee of University College London and complied with all relevant ethical regulations. Before taking part in the experiment, participants were familiarized with the experimental setup and procedures.

### Experimental procedures

Throughout the experiment, participants were seated comfortably in front of a table in a silent, temperature-controlled room, with the palm of their left (non-dominant) hand resting on the table. Each participant received tonic nociceptive stimuli on the dorsum of their left hand, as well as tonic auditory stimuli (see Sensory stimuli, below). In a number of trials of each stimulus type (15 out of 30 high-pain stimuli, 15 out 15 low-pain stimuli, and 15 out of 30 auditory stimuli), participants were instructed to continuously rate their perceived stimulus intensity on a visual analogue scale (VAS) ranging from 0 to 10 using a custom-built vertical slider controlled with their right hand. The slider was connected to a potentiometer to record their ratings. For nociceptive stimuli, the lower and upper ends of the slider were defined as “no pain” and “the maximum pain tolerable” respectively. For the auditory stimuli, they were defined as “no sound” and “the loudest sound tolerable”. Ratings were recorded from the onset of the periodic stimulation, and lasted for 30 s in auditory trials and 45 s in pain trials. Rating data was digitized at 1024 Hz (USB-1408FS, Measurement Computing Corporation, Norton, MA, USA) and synchronized with stimulation triggers and EEG recordings. In all trials, participants were instructed to focus their attention on the stimuli and keep their gaze on a fixation cross placed centrally in front of them, at a distance of approximately 60 cm and 30° below eye level. The order of stimulus presentation and rating task is detailed in **Table S2**. The experimenters started trials manually after ensuring that the participant was ready and instructed whether to rate the sensation. As a result, the time between the end of a trial and the beginning of the following trial ranged between 10.0 and 78.2 s (average 21.2 s). Participants were allowed to rest for approximately 2 minutes after every 15 trials.

### Sensory stimuli

Nociceptive tonic stimuli were generated by a temperature-controlled CO_2_ laser stimulator (Laser Stimulation Device, SIFEC, Ferrières, Belgium) with a wavelength of 10.6 μm and a beam diameter of 6 mm. The output power of the laser was continuously regulated by a feedback control loop based on an online monitoring of skin temperature at the target site (67). The laser stimulation target was changed after each stimulus to avoid nociceptor fatigue, sensitization, and skin damage. Laser power modulation resulted in a 0.1-Hz sinusoidal modulation of skin temperature, starting with an initial phase of π (i.e., trough), and lasting for 30 s (**Fig 1A**). The temperature difference between peaks and troughs was 3°C. Each participant received laser stimuli of two intensity levels (high-pain and low-pain stimuli), individually adjusted as described below. To avoid saliency-related brain responses during the periodic stimulation, the skin temperature was first brought to the desired trough level in a 1-s heating ramp and then maintained at this level for 5 s before the onset of the periodic stimulation. After the periodic stimulation, the skin temperature was maintained at the trough level for 10 additional seconds.

Before the experiment, stimulus temperatures were determined individually as follows. First, pain detection threshold and pain tolerance were estimated. To measure the pain detection threshold, participants received linearly increasing stimuli at 1°C/s (with a cutoff at 54°C) on the dorsum of their left hand, and were instructed to press a button with their right hand as soon as they felt a painful sensation. The button-press immediately terminated the stimulation. To measure the pain tolerance, participants were instructed to press the button as soon as they felt the painful sensation become intolerable. The laser target was changed after each stimulus. Both the pain detection threshold and pain tolerance were measured three times, in consecutive trials, and their corresponding temperatures were estimated as the mean of the three consecutive measurements. The trough temperature of the low-pain stimuli was set to 1°C above the pain detection threshold, and the trough temperature of the high-pain stimuli was set to 1°C above that of the low-pain stimuli. The peak temperature of the stimulus was always below the pain tolerance. The resulting peak temperature in the high-pain stimuli was 48.3 ± 1.9°C (mean ± SD across participants).

Auditory stimuli were generated by MATLAB (MathWorks, Natick, MA, USA) and presented through headphones binaurally. Auditory stimuli consisted of a pure tone (frequency of 280 Hz) whose amplitude was sinusoidally modulated at 0.1 Hz for 30 s (**Fig 1B**). As for the nociceptive stimulation, the sinusoidal modulation started with an initial phase of π (i.e., trough). The sound intensity was individually adjusted to ensure that perceived intensity was similar to the high-pain condition. Auditory stimuli were presented using the Psychophysics Toolbox (68).

### Analysis of subjective sensations

Single-trial rating time series were down-sampled to 512 Hz and smoothed using a moving mean filter with a 1-s window. Filtered data was averaged across trials, for each subject and condition. The peak latency and amplitude of each rating cycle were measured from the average waveforms. The rating peak latencies were measured with respect to the corresponding peak latencies in the stimulus time series (i.e., 5, 15, and 25 s; **Fig 1**). A two-way repeated-measures ANOVA with factors Condition (three levels: high pain, low pain, and sound) and Cycle (three levels: cycle 1-3) was performed separately for the peak latency and amplitude. Post hoc paired-sample two-tailed *t*-tests were performed when a significant (*P*<0.05) main effect or interaction was found (False Discovery Rate [FDR] corrected; the same for post hoc tests described throughout the text).

### EEG recording and preprocessing

EEG was recorded continuously using 128 Ag/AgCl electrodes (SD-128, Micromed S.p.A., Treviso, Italy) placed on the scalp according to the 10-5 system. The EEG signal was sampled at 512 Hz, referenced to the nose, high-pass filtered at 0.02 Hz, with a ground at electrode AFz. Electrooculographic signals were simultaneously recorded using two surface Ag/AgCl electrodes, one placed below the lower eyelid and one laterally to the outer canthus of the right eye. Electrode impedances were kept below 10 kΩ.

EEG preprocessing was conducted using the EEGLAB toolbox (69) and custom-written MATLAB scripts. Continuous EEG data was notch filtered at 50 Hz and harmonics to remove power line noise, and then segmented into epochs ranging from −10 s to 45 s relative to the onset of the sinusoidal stimulation. Differences in EEG baseline across trials were removed by demeaning each trial. The EEG data was re-referenced to the common average. Signals contaminated by eye blinks, eye movements, or muscle activities were corrected using independent component analysis (70). Trials containing excessive signal fluctuations in at least one electrode (amplitude exceeding ± 500 μV) were excluded from further analyses. These trials constituted 4.6% of the total number of trials. The corresponding trials in the rating data were also excluded.

Both frequency and phase measures of entrained neural oscillations can be confounded by transient “evoked-type” responses that repeat at the stimulus frequency (6, 7, 41, 44–47). We did observe a transient response (lasting ~0.5 to 2 s) locked to the increase of auditory stimulation intensity (**Fig S4**). Although interesting, this response could contaminate our measures of ultra-low frequency neural oscillations at the stimulus frequency. We therefore applied a cascade of filters at specifically-defined scales in the time domain to both pain and auditory trials, to minimize the potential confounding effects of such regularly occurring transient responses. We first denoised single-trial data after the above processing steps (*s0*) using a moving mean filter with a 0.5-s window (*s1*). We then applied a 2-s median filter and a Gaussian filter with full-width at half-maximum (FWHM) of 1 s to *s1*, yielding a signal (*s2*) in which the transient responses were removed while the long-period signals were kept. Finally, we reconstructed the EEG signal (*s’*, without the transient responses) as *s’* = *s0* − *s1* + *s2*. This algorithm was effective in removing the transient responses while leaving other features of the signal largely intact (**Fig S4**). Thus, the power increase and phase locking of the EEG responses revealed by the following analyses were most likely due to a true 0.1-Hz oscillation rather than transient responses that repeated at this frequency.

### Power analysis of EEG data

A fast Fourier transform was applied to single-trial signals ranging from 0 to 30 s after the onset of the sinusoidal stimulation, yielding power spectra with a frequency resolution of 0.0333 Hz (a frequency resolution of 0.01 Hz, i.e., spectral interpolation, was achieved by zero padding in the time domain and was used for illustrative purposes). Power estimates were log-transformed and averaged across trials for each subject and condition. To reveal the frequency of power increases, for each subject, condition, electrode and frequency point, the contribution of background activities (e.g., spontaneous brain activities or slow eye movements) was removed by subtracting the average power at surrounding frequencies (−0.0333 Hz and +0.0333 Hz) (20, 36, 37). Scalp topographies of this background-subtracted power (BSP) were computed by spline interpolation.

To identify the frequencies at which power increase occurred, a one-sample one-tailed *t*-test was performed at each frequency point to test whether the BSP was consistently greater than zero *across subjects* (FDR corrected for multiple comparisons across frequencies). This analysis was first performed on signals from a central electrode cluster (Cz and its closest neighbours FCC1h, FCC2h, CCP1h, and CCP2h) and then extended to all electrodes (FDR corrected for multiple comparisons across frequencies and electrodes). An additional one-sample one-tailed *t*-test was performed separately for each subject and condition, to examine whether the BSP at 0.1 Hz in the central electrode cluster was consistently greater than zero *across single trials*.

To test the effects of modality and rating task on the power increase detected at 0.1 Hz, we conducted, for each electrode, a two-way repeated-measures ANOVA with factors Modality (two levels: high pain and sound) and Rating (two levels: rating and no rating) on 0.1-Hz BSP (FDR corrected for multiple comparisons across electrodes). Post hoc paired-sample two-tailed *t*-tests were performed when a significant main effect or interaction was found.

### Phase analysis of EEG data

We examined the phase locking of the EEG signal (0-30 s) across trials by calculating the inter-trial phase coherence (ITPC) (71) for each subject, condition, and electrode. Briefly, given the Fourier phase *φ*_*n*_ for trial n, we define the mean vector of phase angles across trials as 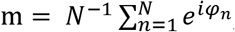, where N is the number of trials. The ITPC value is given by the modulus of m, i.e., ITPC = |m|. To determine at which frequencies phase locking occurred, we evaluated the ITPC as a function of frequency by calculating the ITPC values in steps of 0.0333 Hz (a frequency resolution of 0.01 Hz was achieved as described in the previous paragraph). ITPC scalp topographies were computed by spline interpolation. For each subject, the significance of ITPC was determined using the Rayleigh’s test for circular uniformity (*P*<0.05) (39). The percentage of subjects with significant ITPC was calculated for each frequency point, electrode, and condition.

To further assess the significance of the ITPC at the group level (i.e., whether ITPC was greater than what one would expect by chance), the mean ITPC across subjects and the percentage of subjects with significant ITPC were compared to randomized data. Specifically, we added random phase values drawn from circular uniform distribution to the single trial phases, recalculated the ITPC for each subject, and determined its significance using the Rayleigh’s test. We then computed the mean ITPC across subjects and the percentage of subjects with significant ITPC. This process was repeated 1,000 times, yielding null distributions of the mean ITPC and of the percentage of subjects with significant ITPC. *P* values of the actual data were determined by comparing the mean ITPC and percentage of subjects to the respective null distributions. This analysis was also first conducted for the 0.1-Hz oscillation in the central electrode cluster, and then extended to other frequencies and electrodes (FDR corrected for multiple comparisons across frequencies and electrodes).

To test the effects of modality and rating on the ITPC at 0.1 Hz, for each electrode, we performed a two-way repeated-measures ANOVA with factors Modality (two levels: high pain and sound) and Rating (two levels: rating and no rating) (FDR corrected for multiple comparisons across electrodes). Post hoc paired-sample two-tailed *t*-tests were performed when a significant main effect or interaction was found.

To examine the dependence of the degree of phase locking on the phase of oscillations occurring before stimulus onset, we applied a causal, linear-phase bandpass FIR filter with cutoff frequencies at 0.05 and 0.15 Hz to single-trial EEG signals from the central electrode cluster, and then extracted the instantaneous phase of the filtered signals using the Hilbert transform. The causal filter was used to avoid the influence of signals after stimulus onset on the prestimulus phase. We sorted the single trials into six bins from 0 to 2π according to the instantaneous phase at time 0 s in the filtered signal. Within each subject, we pooled rating and no-rating trials from the same modality, to increase the number of trials in each bin. We then calculated 0.1-Hz ITPC during the sinusoidal stimulation (0-30 s) for the trials within each bin, separately for each subject and modality. Since the number of trials influences ITPC (i.e., fewer trials are more likely to have a greater ITPC value) (72), to correct for differences in the number of trials between bins and modalities, we transformed the ITPC to ITPCz (Rayleigh’s Z) according to the formula *ITPC*_*z*_ = *N* · *ITPC*^2^, where N is the number of trials for each ITPC calculation, as previously recommended (73–75). Since the causal filter introduced a delay in the filtered signal, we shifted the phase bins accordingly (by ½ cycle of a 0.1-Hz oscillation) to estimate the relationship between the ITPC_z_ and the instantaneous phase at the onset of the rhythmic stimulation (importantly, this was not a shift of the filtered signal and did not introduce non-causality). We performed a two-way repeated-measures ANOVA on the ITPCz with factors Bin (six levels: equal-sized bins from 0 to 2π) and Modality (two levels: pain and sound). Post hoc paired-sample two-tailed *t*-tests were performed when a significant main effect or interaction was found.

### Analysis of relationship between the strength of neural entrainment and perceived pain intensity across individuals

For each condition entailing a rating task, we calculated across-subject Pearson correlation coefficients between the intensity rating at each time point in the rating time series and the 0.1-Hz BSP as well as the 0.1-Hz ITPC in the central electrode cluster, yielding time series of the correlation coefficient *r* and the *P* value (FDR corrected for multiple comparisons across time points). To test whether the correlations were consequent to the rating task, we performed the same correlation analyses between the intensity ratings and the BSP/ITPC measured in the conditions without rating task. Finally, the same correlation analyses were also performed between conditions entailing different intensities of nociceptive stimulation.

For each subject and condition, we calculated the phase of the entrained 0.1-Hz oscillations as the orientation of the above-defined mean vector m (i.e., arg(m)). To evaluate the across-subject relationship between the intensity rating and the phase of entrained oscillations, we fitted a single-cycle cosine to the subject-mean peak rating (i.e., the peak rating averaged across the three cycles) as a function of the phase of 0.1-Hz oscillations in the central electrode cluster. Significance of the cosine fit was estimated with permutation testing: we randomly permuted the phase values across subjects, fitted a cosine function, and calculated the coefficient of determination *R*^2^ as a measure of the goodness of fit. This procedure was repeated 1,000 times, yielding null distribution of the *R*^2^. The *P* value of *R*^2^ obtained from the actual data was determined by comparing it to the null distribution. This analysis was performed between intensity rating and 0.1-Hz phase within each condition entailing the rating task, between intensity rating and 0.1-Hz phase in the conditions without rating task, and between conditions entailing different intensities of nociceptive stimulation.

To ensure that the results from the above analyses were not due to individual variability in stimulus temperature, we performed similar analyses but using stimulus temperature instead of ratings, and also analyzed the correlation between stimulus temperature and pain ratings. Thus, we analyzed across-subject relationships between the laser stimulation temperature and (i) mean peak pain rating (i.e., the peak rating averaged across the three cycles), (ii) the BSP, (iii) the ITPC, and (iv) the phase of 0.1-Hz oscillations in the central electrode cluster.

### Analysis of neural oscillations outlasting the stimulus

The EEG time series from each subject was smoothed by a moving mean filter with a 2-s window, and linearly detrended. No zero padding was used at the signal edges, to ensure that any oscillation after stimulus offset was not due to the temporal smoothing. A one-sample two-tailed *t*-test of EEG amplitude against zero was performed at each point of the time series (FDR corrected for multiple comparisons across time points). Scalp topographies of the *t* value were computed over a 1-s window centered around the peak and trough of each cycle. Finally, Pearson correlation coefficients were calculated across subjects between the mean peak rating and the peak-to-trough amplitude of the cycle occurring after the sinusoidal stimulation in the central electrode cluster.

## Supplementary Information

### Supplementary psychophysical results

For each participant and condition, we extracted the peak amplitude (**Fig 1C**, **top**) and latency (**Fig 1C**, **bottom**) of each of the three rating cycles. Two-way repeated-measures ANOVAs with factors Condition (three levels: high pain, low pain, and sound) and Cycle (three levels: 1-3) revealed strong evidence for main effects of the two factors and their interaction (Peak amplitude: Condition, *F*_2,58_=43.09, *P*<0.0001, partial *η*^2^=0.5977; Cycle, *F*_2,58_=86.79, *P*<0.0001, partial *η*^2^=0.7496; interaction, *F*_4,116_=67.62, *P*<0.0001, partial *η*^2^=0.6999. Peak latency: Condition, *F*_2,58_=72.85, *P*<0.0001, partial *η*^2^=0.7153; Cycle, *F*_2,58_=58.61, *P*<0.0001, partial *η*^2^=0.6690; interaction, *F*_4,116_=26.70, *P*<0.0001, partial *η*^2^=0.4793). The peak rating amplitude was higher in the high pain than in the low pain condition (post hoc tests: *P*<0.0001 for each of the three cycles), higher in the auditory than in the low pain condition (*P*<0.0001 for each of the three cycles), as well as higher in the auditory than in the high pain condition, in the last two cycles (both *P*<0.0001) but not in the first cycle (*P*=0.1455). Furthermore, auditory ratings peaked earlier than pain ratings in all three cycles (auditory vs. high pain: *P*=0.0151 [cycle 1], *P*<0.0001 [cycles 2-3]; auditory vs. low pain, *P*<0.0001 [all cycles]), whereas peak latencies in the two pain conditions were not significantly different except for the first cycle, in which high pain latencies were shorter than low pain latencies (cycle 1: *P*=0.0023; cycle 2: *P*=0.7670; cycle 3: *P*=0.0791). Finally, peak amplitudes of pain ratings were smaller and delayed in the last two cycles compared to the first cycle (amplitude: in both high and low pain, cycle 1 vs. cycle 2 or 3, all *P*<0.0001; latency: in both high and low pain, cycle 1 vs. cycle 2 or 3, all *P*<0.0001). This was not the case for the auditory ratings (*P*>0.2009 in all comparisons).

**Table S1.** Across-subjects relationship between stimulation temperature and pain ratings, as well as between stimulation temperature and different features of neural entrainment at 0.1 Hz.

|  | High Pain, No Rating | High Pain, Rating | Low Pain, Rating |
| --- | --- | --- | --- |
| **Pain intensity rating** | | $r = -0.2015, P = 0.2857$ | $r = -0.1206, P = 0.5256$ |
| **BSP** | $r = -0.2142, P = 0.2557$ | $r = 0.1955, P = 0.3005$ | $r = -0.0715, P = 0.7071$ |
| **ITPC** | $r = -0.0472, P = 0.8042$ | $r = 0.0962, P = 0.6130$ | $r = 0.0020, P = 0.9916$ |
| **Phase** | $\rho_{cl} = 0.2577, P = 0.3695$ | $\rho_{cl} = 0.1566, P = 0.6922$ | $\rho_{cl} = 0.0466, P = 0.9679$ |

BSP: background-subtracted power. ITPC: inter-trial phase coherence. Pain ratings were the peak amplitude averaged across three cycles, in each subject. The 0.1-Hz BSP, ITPC, and phase were measured from the central electrode cluster. For pain ratings, BSP and ITPC, the relationships are expressed as Pearson's correlation. For Phase, the relationship is expressed as circular-linear correlation<sup>1</sup>. N=30 subjects.
<sup>1</sup> P. Berens, CircStat: a MATLAB toolbox for circular statistics. *J Stat Softw* **31**, 1-21 (2009).

**Table S2.** Order of stimulation trials.

|  |  |  |  |  |  |  |  |  |  |  |  |  |  |  |  |
| --- | --- | --- | --- | --- | --- | --- | --- | --- | --- | --- | --- | --- | --- | --- | --- |
| **Trials**<br>**1-15** | HP<br>R | AUD<br>R | LP<br>R | HP<br>nR | AUD<br>R | LP<br>R | HP<br>R | AUD<br>nR | LP<br>R | HP<br>nR | AUD<br>R | LP<br>R | HP<br>R | AUD<br>nR | LP<br>R |
| **Trials**<br>**16-30** | HP<br>nR | AUD<br>R | LP<br>R | HP<br>R | AUD<br>nR | LP<br>R | HP<br>nR | AUD<br>R | LP<br>R | HP<br>R | AUD<br>nR | LP<br>R | HP<br>nR | AUD<br>R | LP<br>R |
| **Trials**<br>**31-45** | HP<br>R | AUD<br>nR | LP<br>R | HP<br>nR | AUD<br>R | LP<br>R | HP<br>R | AUD<br>nR | LP<br>R | HP<br>nR | AUD<br>R | LP<br>R | HP<br>R | AUD<br>nR | LP<br>R |
| **Trials**<br>**46-60** | HP<br>nR | AUD<br>R | HP<br>R | AUD<br>nR | HP<br>nR | AUD<br>R | HP<br>R | AUD<br>nR | HP<br>nR | AUD<br>R | HP<br>R | AUD<br>nR | HP<br>nR | AUD<br>R | HP<br>R |
| **Trials**<br>**61-75** | AUD<br>nR | HP<br>nR | AUD<br>R | HP<br>R | AUD<br>nR | HP<br>nR | AUD<br>R | HP<br>R | AUD<br>nR | HP<br>nR | AUD<br>R | HP<br>R | AUD<br>nR | HP<br>nR | AUD<br>nR |

HP: high-pain stimulation. LP: low-pain stimulation. AUD: auditory stimulation. R: trials in which participants were required to continuously rate perceived intensity. nR: trials without the rating task. Participants were allowed to rest for approximately 2 minutes after every 15 trials.

**Figure S1.**
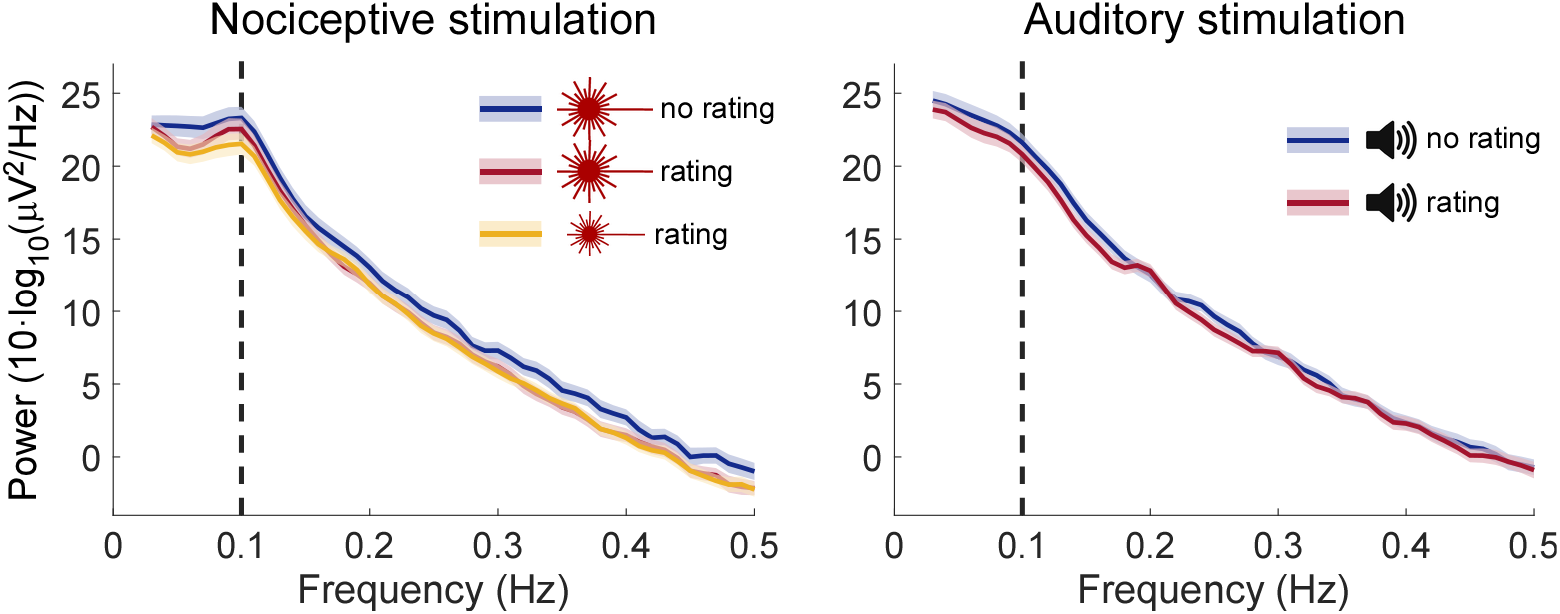
Absolute EEG spectra during the rhythmic nociceptive (left) and auditory (right) stimulation in the central electrode cluster. The central electrode cluster included Cz and its closest neighbours FCC1h, FCC2h, CCP1h, and CCP2h. Shaded regions indicate SEM across subjects (N=30). The vertical dashed line indicates the frequency of stimulation.

**Figure S2.**
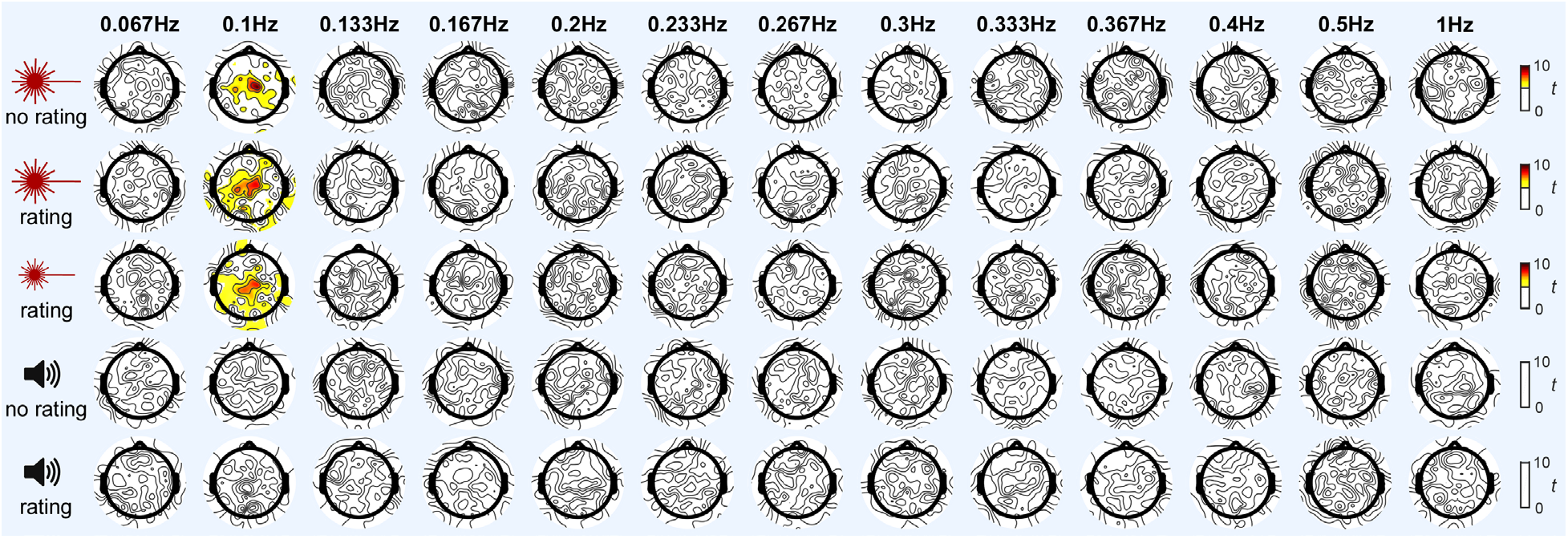
Topographies of EEG power enhancement at different frequencies. Topographies of *t* values show strong evidence of EEG power enhancement (expressed as background-subtracted power, BSP) at 0.1 Hz in central scalp regions, only in the conditions with nociceptive stimulation. Colours indicate scalp electrodes where the BSP had *P*<0.05 (one-sample *t*-test against 0, FDR corrected across electrodes and frequencies). Electrodes with *P*>0.05 are masked with white. N=30 subjects.

**Figure S3.**
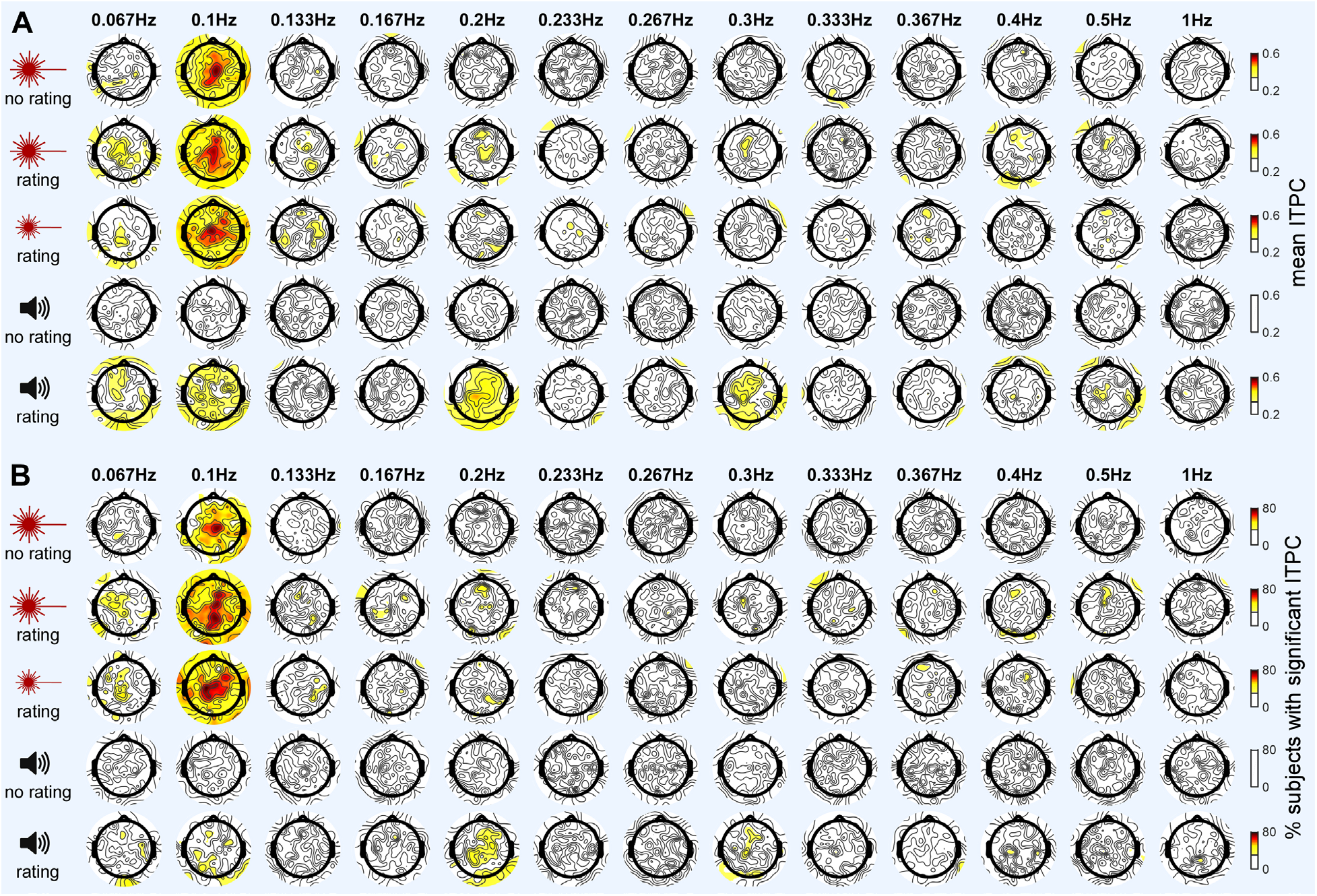
Topographies of EEG phase locking at different frequencies. Topographies showing strong evidence of EEG phase locking at 0.1 Hz in central scalp regions (using two measures: mean ITPC, panel **A**; percentage of subjects with significant ITPC, panel **B**), mostly in the conditions with nociceptive stimulation. There was a weak suggestion of phase locking at 0.2 and 0.3 Hz in the auditory condition which also entailed rating. Colours indicate scalp electrodes where the phase locking was greater than chance level (comparison with randomized data; *P*<0.05, FDR corrected across electrodes and frequencies). Electrodes with *P*>0.05 are masked with white. N=30 subjects.

**Figure S4.**
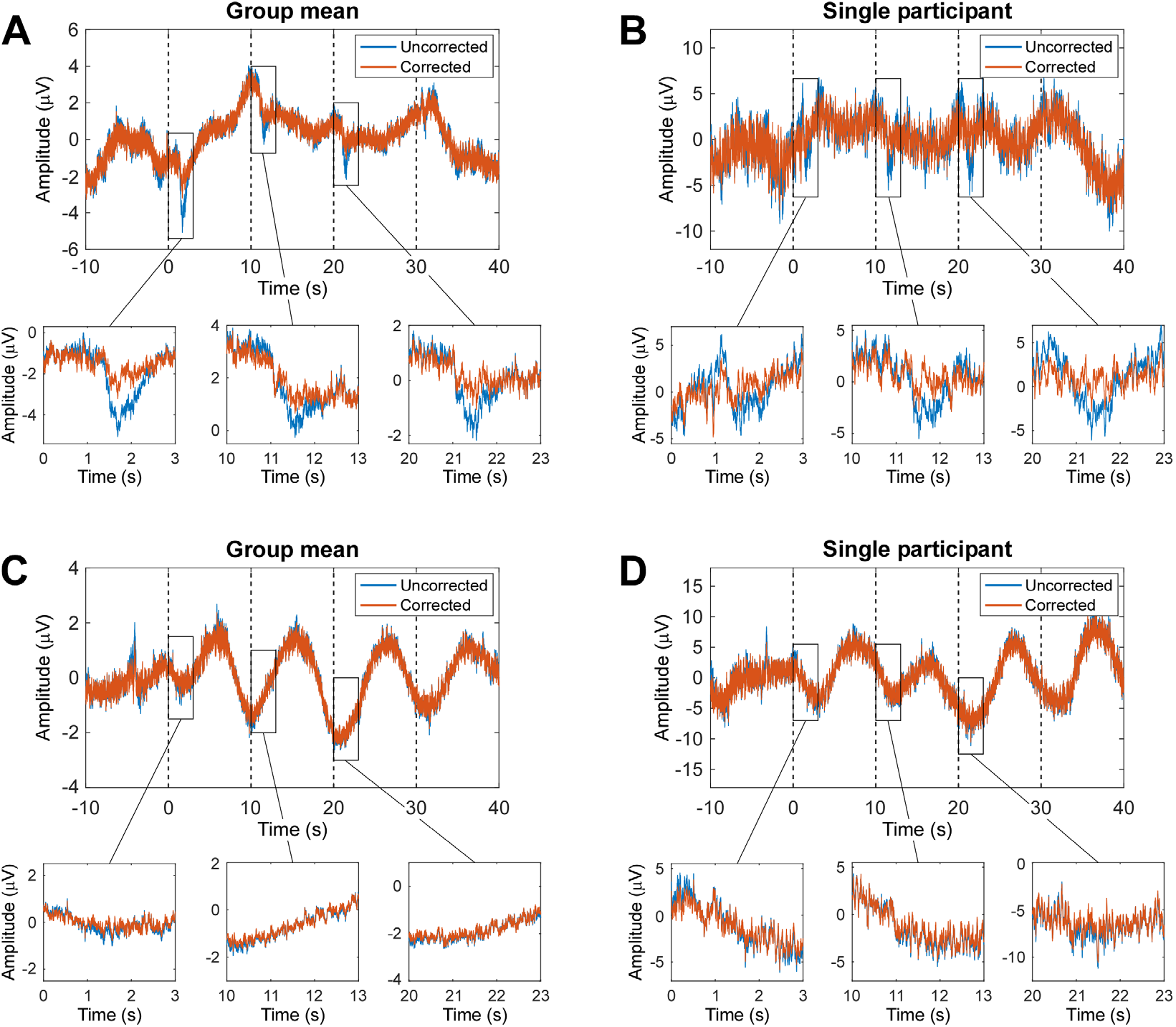
Minimizing the confounding effect of transient responses observed during auditory stimulation with rating. (**A**) EEG signal recorded at electrode Fz (the electrode showing the largest transient response) during auditory stimulation with rating condition, averaged across subjects (N=30). Transient responses are visible during each of the three increases of stimulus intensity (blue line, uncorrected signal). The correction algorithm (see Materials and Methods) effectively suppressed these transient responses while leaving other features of the signal largely intact (red line, corrected signal). Insets show magnified views of the regions indicated by rectangles. (**B**) Same as (A), but showing signal from a participant with clear transient responses. (**C**) EEG signal recorded at electrode Cz during high-pain with rating condition, averaged across subjects. Note the lack of the transient responses observed in the auditory condition: the correction algorithm barely affected the recorded signal. (**D**) Same as (C), but showing signal from a single participant.

